# Isolation and characterization of a Cyprinid herpesvirus strain YZ01 from apparently healthy goldfish after rising water temperature

**DOI:** 10.1101/2021.07.05.451087

**Authors:** Fengzhi Wang, Ye Xu, Yi Zhou, Chao Ding, Hongan Duan

## Abstract

A Cyprinid herpesvirus 2 (CyHV-2) strain YZ01 was isolated from an apparently healthy goldfish with history of exposure to CyHV-2 when the water temperature was heated from 3-6□ to 25□. Brain, kidney, spleen (referred to as BKS) and intestine of sick and dead fish as well as crucian carp (CrCB) or goldfish brain cell (GFB) suspensions inoculated with above tissue homogenate were collected for PCR assay.Infected cell layers were prepared into ultrathin sections for transmission electron microscopy. Virus DNA were purified from cell suspensions for whole genome sequencing and analysis. Intestine and mixed brain, kidney and spleen fluid (BKS) of dead goldfish and BKS-infected CrCB suspensions as well as their subsequent passages were tested positive by PCR. Nucleus deformation, lattice-like arrangement of inclusion bodies and various forms of immature and mature virions in the nucleus, cytoplasm and outside of cell membranes were observed. Whole genome analysis of 7 CyHV-2 strains available in GeneBank showed that all strains’ genomes share very high homology while YZ01 was more close to CyHV-2 CNDF-TB2015 and SY-C1 which are classified as C genotype.

## Introduction

Cyprinid herpesvirus-2 (CyHV-2), early known as the cause of herpesviral hematopoietic necrosis disease (HVHND) is highly pathogenic and causes huge losses to goldfish (*Carassius auratus*), crucian carp (*C. carassius*) and Gibel carp (*C. auratus gibelio*). The disease was first discovered in goldfish in Japan during 1992-1993 and has spread all over the world where goldfish and crucian carp is cultured. Detection and research work mainly depend upon the detection of etiologic agents based on the virus morphology and/or virus DNA. Various virus stages and different size ranging from hexagonal nucleocapsids of 95×106 nm(A. E. Goodwin et al., 2006), hexagonal enveloped virus particles of 170-220 nm(Luo et al., 2013), viral nucleocapsids of 95-110 nm and enveloped viral particles of 170-200 nm (Wu et al., 2013), nucleocapsid with 90–120 nm and mature enveloped virus of 170-200 nm, negatively stained purified virions of 110-120 nm(Fichi et al., 2013; J. Xu et al., 2013) had been observed in nucleus, cytoplasm and extracellular spaces of affected tissues. As other herpesviruses CyHV-2 replicates and assembles in hematopoietic cells of spleen, kidney and gills. Mature process includes budding into vesicles inside cytoplasm with envelope containing viral glycoproteins and releasing outside of the cells. The various virion forms and replication process were described in more detail in a comprehensive review(Thangaraj et al., 2020). Genetic homology analysis based on partial nucleotide sequences of DNA polymerase showed that CyHV-2 isolated in USA and China were highly homological to ST-J1 isolated from Japan (A. E. Goodwin et al., 2006; Li et al., 2015), even though some differences exist in predicted amino acid sequences of certain proteins of ST-J1 and strains from China. Researches also revealed different lengths of Marker A(mA) region of viral genome of several CyHV-2 strains and suggested CyHV-2 be divided into pleural genotypes (Boitard et al., 2015; Ito et al., 2017). Later whole genome comparison of CyHV-2 strains ST-J1, SY-C1 (Davison et al., 2012; Li et al., 2015), CaHV (Zeng et al., 2016), SY(Liu et al., 2018) showed that different strains’ genome had mutations, insertions, deletions and rearrangements although all strains’ genomes were about 98% homological (Li et al., 2015; Liu et al., 2018). These strains were suggested to be divided into C (China genotype) and J genotype (Japan genotype) and main genotype of CyHV-2 in China is closer to C genotype than J genotype (Li et al., 2015). A report discovered variations among genomes of CyHV-2 strains newly isolated from crucian carp in China and C and J genotypes(Liu et al., 2018). In this study a CyHV-2 stain YZ01 (available in GeneBank as YZ01, accession number MK260012.1 hereafter YZ01) was isolated from apparently healthy crucian carp in a farm in Jiangsu Province after raised water temperature. Electron microscopy examination of YZ01 infected GFB cell line and whole genome comparison with those published in GeneBank were carried out in order to further investigate characteristics and epidemiological features of this isolate.

## Materials and methods

### Experimental infection of goldfish

Clinically healthy goldfish were collected from ornamental fish farms with a history of CyHV-2 infection. Goldfish were sampled on February 7, 2017 and delivered to the laboratory on the 9th and placed in tanks. The fish were approximately 8-10 cm in length and had no obvious clinical signs. After acclimatization to the laboratory environment, the tank water was heated to approximately 25°C. If dead fish were present, the brain, spleen, kidney mixture and intestine of the dead fish were taken separately, added to MEM medium at a volume ratio of 1:10, and a portion of tissue homogenized was taken for nucleic acid extraction and PCR detection. CyHV-2 PCR assay was performed using the following primers:CyHV-2pol-FOR(CCCAGCAACATGTGCGACGG), CyHV-2pol-REV (CCGTARTGAGAGAGTTGGCGCA) with Fragment size of 362bp. The remaining homogenate for virus isolation on cell culture was added to mixture of streptomycin and penicillin, overnight at 4°C. The PCR-positive homogenate was selected, centrifuged at 4000×g for 15 min, the supernatant was filtered through a 0.45μm sterile filter, and 1 mL/bottle was inoculated onto crucian carp brain cells (CrCB) or goldfish brain cells (GFB) (Y. Xu et al., 2019) over 80% fluent monolayer. Cells were observed daily for growth status and CPE. If cells were over 80% CPE, the suspension was transferred to the sub-passage.

### Preparation of ultrathin sections for electron microscopy observation

Virus-infected cell samples and ultra-thin sections were prepared similarly as reported(Dong et al., 2011). Briefly the 14th passage(P14) of CyHV-2 isolate YZ01 was inoculated onto GFB cell lines grown in 3 culture flasks (25m^2^, Corning 430168). The medium was poured out 6 days after inoculation (dpi) when over 60% CPE appeared, the cells were scraped off, collected to centrifuge at 15000 rpm for 10 min to make rice grains large, and then pre-fixed in 0.1 M phosphate buffer containing 2.5% glutaraldehyde and fixed in 0.1 M phosphate buffer containing 2.0% osmium tetroxide. Ultrathin sections were stained with uranyl acetate-lead citrate and examined with a TEM (H-7650, 1600CCD)Hitachi transmission electron microscope (Shanghai Chenmai New Material Testing Center).

### Whole genome sequencing and analysis

The 13th passage of CyHV-2 isolate strain YZ01 was inoculated onto GFB cell lines. When complete CPE was present, approximately 20 mL of cell suspension was centrifuged at 7,000 × g for 10 min, followed by ultra-centrifugation at 120,000 × g for 1 hr. Pellet was re-suspended in 1 mL PBS and sent to Shanghai Hanyu Biotechnology Co., Ltd. for sequencing. The genome sequence was then submitted to GeneBank. The accession number was MK260012.1, named as CyHV-2 strain YZ01. The entire genome of YZ01 and all other CyHV-2 isolates available in GeneBank (Table 1) was subjected to multiple comparisons and phylogenic tree construction using MEGA X software(Kumar et al., 2018)and PhyML online execution by default options available on website http://www.atgc-montpellier.fr/phyml/. Phylogenic relationship was inferred using the unweighted pair group method with arithmetic mean method(UPGMA), neighbor-joining(NJ) and maximum evolution method(ME) by MEGA X. The bootstrap consensus tree inferred from 1000 replicates was taken to represent the evolutionary history of the strains analyzed. Branches corresponding to partitions reproduced in less than 50% bootstrap replicates were collapsed. The percentage of replicate trees in which the associated strains clustered together in the bootstrap test (1000) was shown next to the branches. The evolutionary distances were computed using the number of differences method and were in the units of the number of base differences per sequence. Codon positions included were 1st+2nd+3rd+Noncoding. All positions containing gaps and missing data were eliminated.

**Table 1:** Whole genomes of CyHV-2 isolates available in GeneBank

| CyHV-2 stains | Length(bp) | Submitted date | Country/area isolated |
| --- | --- | --- | --- |
| NC_019495 (JQ815364) | 290304 | Nov21 2012 | Japan |
| CyHV-2 ST-J1 |  |  |  |
| KM200722CyHV-2SY-C1 | 289365 | Jul 15 2014 | China |
| KT387800 CyHV-2 SY | 290455 | Aug12 2015 | China |
| KU199244 CyHV-2 1301 | 275348 | Nov27 2015 | China |
| MK260012 CyHV-2 YZ01 | 288076 | Dec02 2018 | China |
| MN201961CyHV-2 | 288321 | Jul 20 2019 | China |
| CNDF-TB2015 |  |  |  |
| MN593216 CyHV-2 YC-01 | 275367 | Oct14 2019 | China |

## Results

The experimental fish fell sick and began to die about a week after the water temperature was heated from 3-6°C to 25°C. Cloudy eyeballs, detaching fish scales and hemorrhage around the mouth was observed in diseased fish(Figure 1). Brain and viscera had no obvious histo-pathological changes. Some fish died without apparently clinical signs. 4 days after inoculation of brain, kidney and spleen (BKS) suspension of the dead goldfish in tank 4 onto CrCB cell layer, it was seen that cells in some focal area became brighter and round. The inter-cell spacing increased and cell boundary became more clear. The focal lesions were obvious. The cells became round and piled up, forming a fibrous network. After 10 days the cell layer was sieve-shaped and clustered (Figure 2).

**Figure 1.**
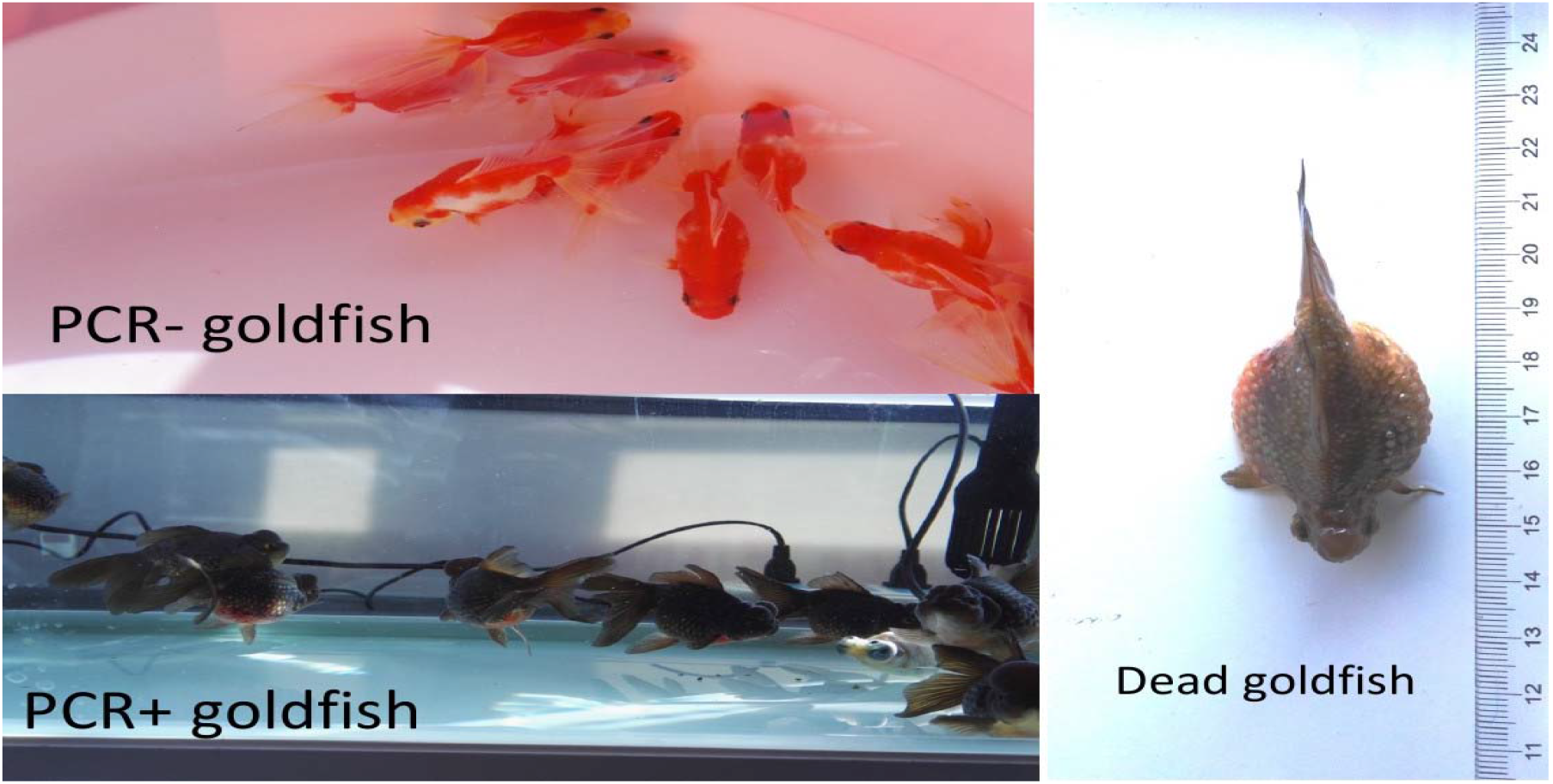
Experimental infection and isolation of CyHV-2 from apparently healthy goldfish

**Figure 2.**
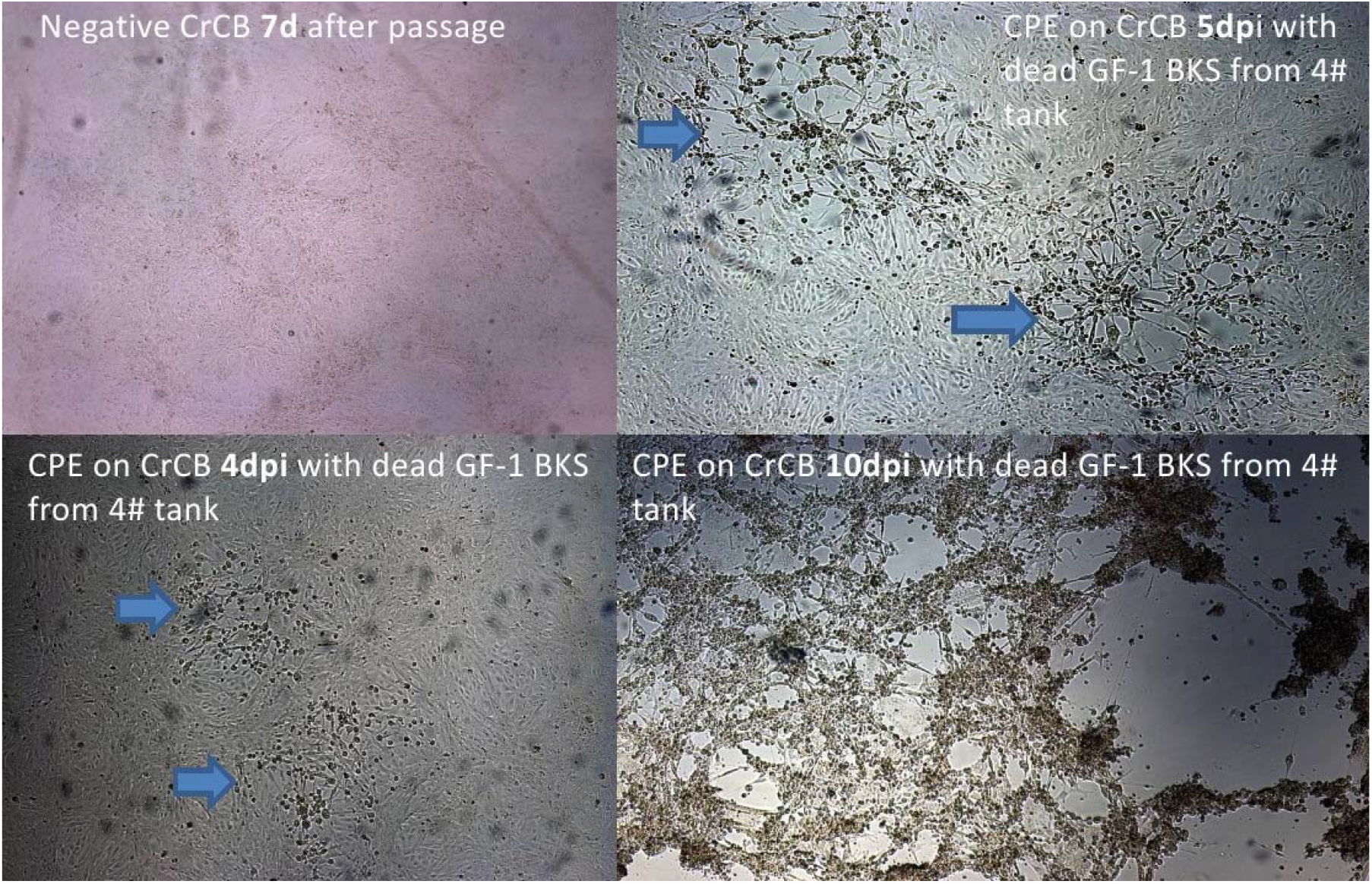
CPE on CrCB inoculated with tissue suspension of dead goldfish. 4 days after inoculation(4 dpi) into CrCB with brain, kidney and spleen(BKS) suspension of the dead goldfish in No. 4 water tank, cell layer showed brightening and rounding in focal areas, and the inter-cell spacing increased; after 5 days, the focal lesions were obvious. The cells became round and piled up, forming a fibrous network. After 10 days, the cell layer was sieve-shaped and clustered.

PCR results were positive for intestine and mixed brain, kidney and spleen fluid (BKS) of dead goldfish and BKS-infected CrCB suspensions (Table 2-3). Subsequent passages were also positive by PCR (Figure 3-4). While sometimes intestine samples or intestine inoculated CrCB suspension tested PCR negative.

**Table 2:** Death of fish after rising water temperature

| 3#tank | Total 13 |  | 4#tank | total 12 |  |
| --- | --- | --- | --- | --- | --- |
| date | death | PCR | death | death | PCR |
| Feb15 | 2 | + | Feb18 | 1 | + |
| Feb17 | 1morbid | + | Feb21 | 1 | + |
| Feb18 | 1 | + | Feb22 | 2 | + |
| Mar8 | 1 | + | Feb25 | 1 | + |
| Mar24 | 1 | + | Feb26 | 1 | + |
|  |  |  | Mar13 | 1 | + |
| total | 7 |  | total | 7 |  |

**Table 3:**
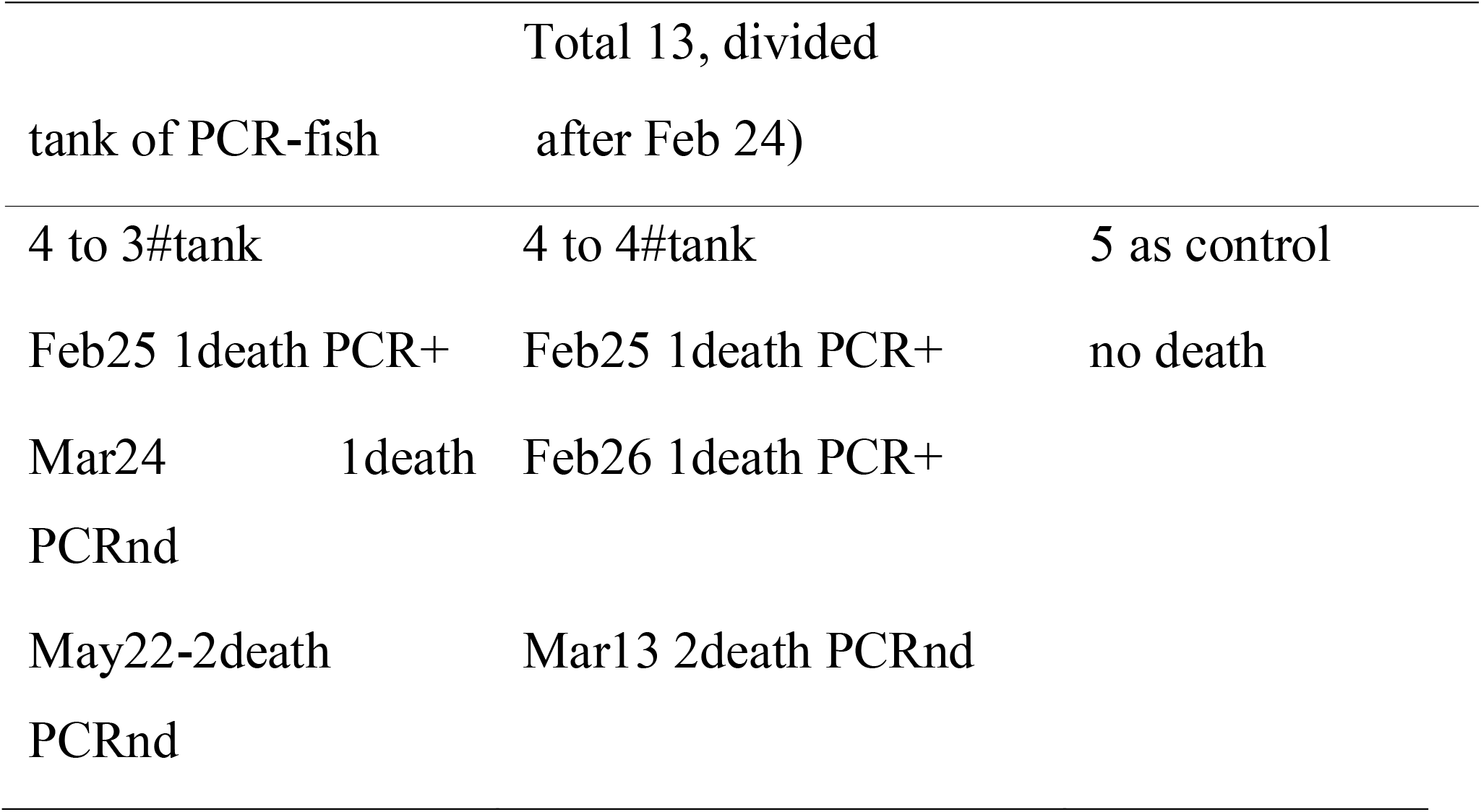
Co-habitation of PCR- fish in 3# and 4# tank

**Figure 3.**
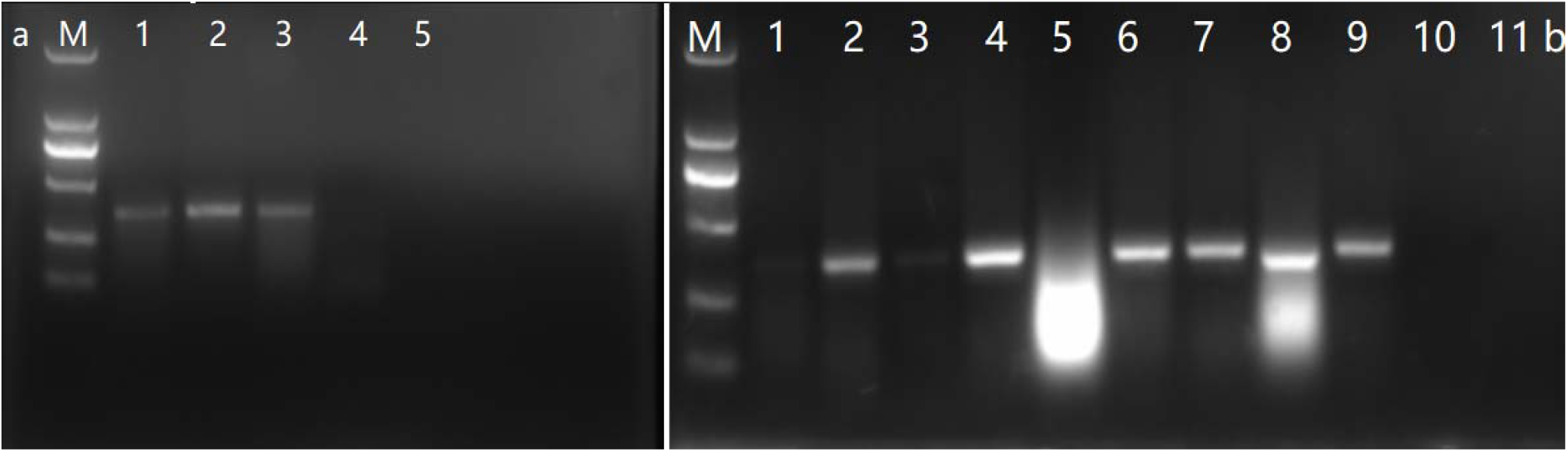
a-PCR results for dead goldfish on Feb 15, 2017. M: DL2000 ladder; lane1: intestine; lane 2: BKS mixture; lane 3: positive control; lane 4: blank control. b-PCR results for CrCB suspension inoculated with infected tissue homogenate. lane1: GF from Xuzhou; lane 2-6: BKS, intestine of DGF on Feb17 and intestine, BKS, brain of DGF on Feb18 in 3#tank. lane 7-8: intestine and BKS of DGF on Feb18 in 4#tank. lane9-11: positive, negative and blank control. Note: DGF: dead goldfish, BKS: brain, kidney and spleen.

**Figure 4.**
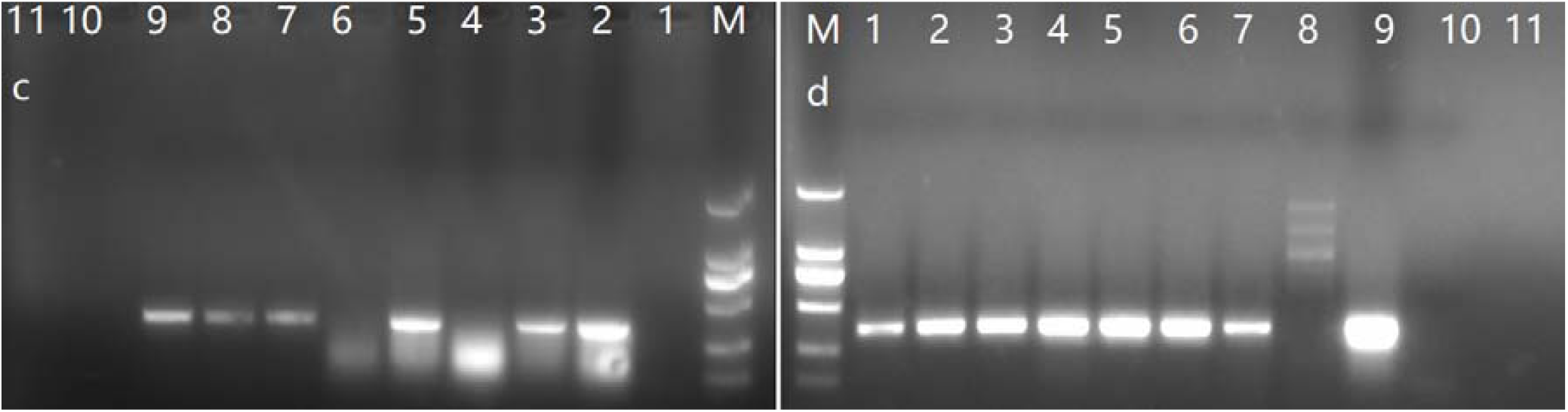
c—PCR results for 2 dead goldfish from 4#tank. M: DL2000; Lane 1-2: blank control, positive control, ;lane3-4: BKS mixture and intestine from No.2 dead goldfish; lane5-6: BKS mixture and intestine from No.1 dead goldfish,lane7-9:CrCB suspension infected on Feb 9,Jan15 and Feb 4 of tissue homogenates of dead fish from Xuzhou; d-PCR results for CrCB suspension. M: DL2000; lane1-8: CrCB suspension infected by BKS of DGF from 4#tank died and inoculated on the CrCB on Feb21(lane1,2),Feb22(lane3,4),Feb25(lane5,6),Feb26(only brain, lane7,8); lane9:positive control; lane10: Blank control; BKS:brain kidney and spleen mix; DGF: dead goldfish.

Electron microscope examination of infected cell samples with tissue homogenate of dead fish showed nucleus deformation, incomplete and invisible nucleus membrane, degenerated or destroyed organelles, lattice-like arrangement of inclusion bodies(figure 5), large number of immature viruses and mature viral particles in the nucleus and cytoplasma or outside of cells(Figure 6-7). 3 forms of nuclocapsids included electron dense, double-layered ring shape and electron lucent nucleocapsid, respectively(Figure 6).Virus in vesicules in cytoplasm, a virus inside the vesicles protruding outside the nuclear membrane were observed(Figure 5-6).

**Figure 5.**
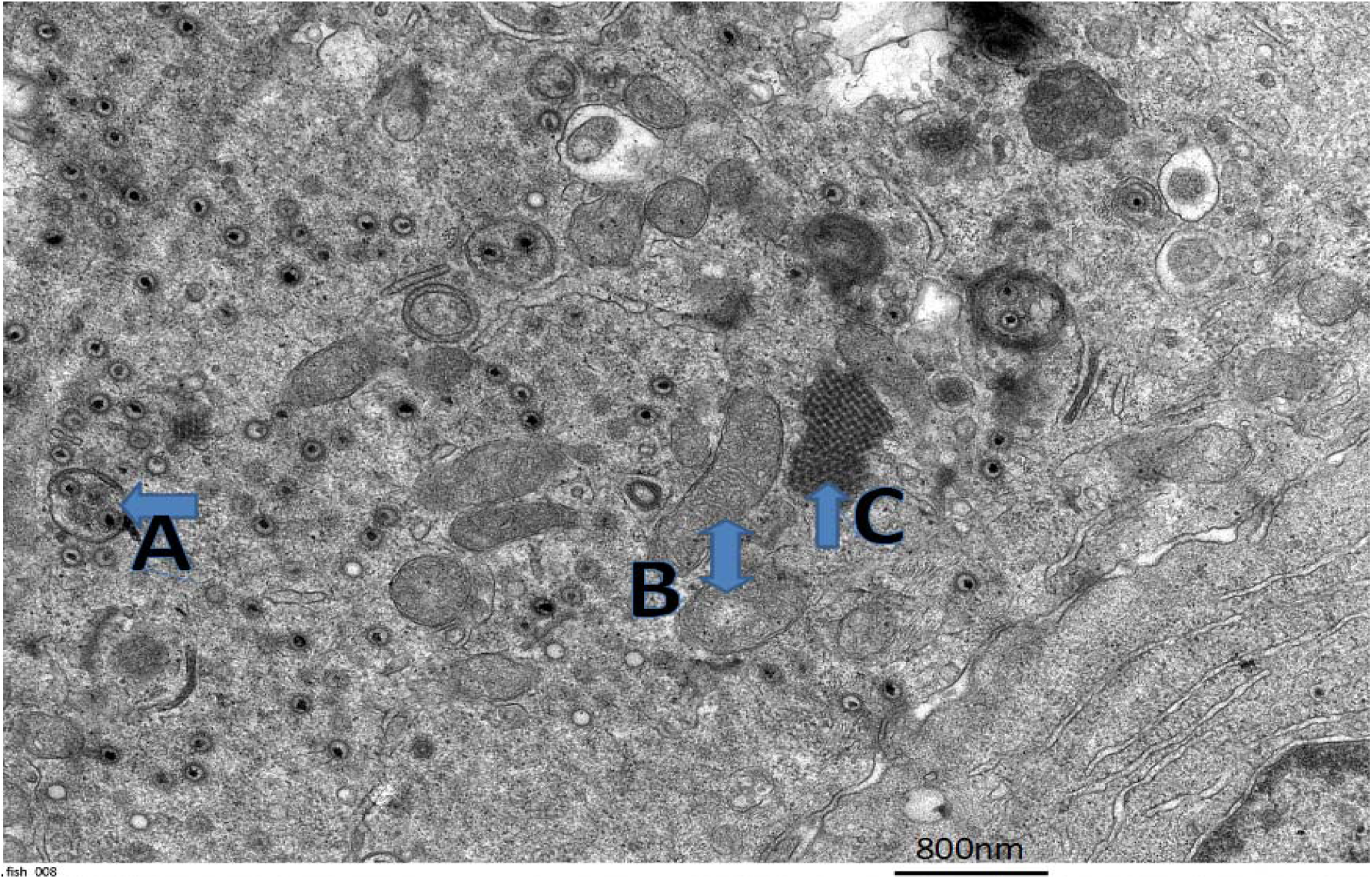
Electron micrographs of goldfish cells(GFB) infected with CyHV-2. The nuclear membrane of the cell is incomplete and invisible. A: virus in vesicules; B: degenerated or destroyed organelles; C: Lattice-like arrangement of nucleocapsids.

**Figure 6.**
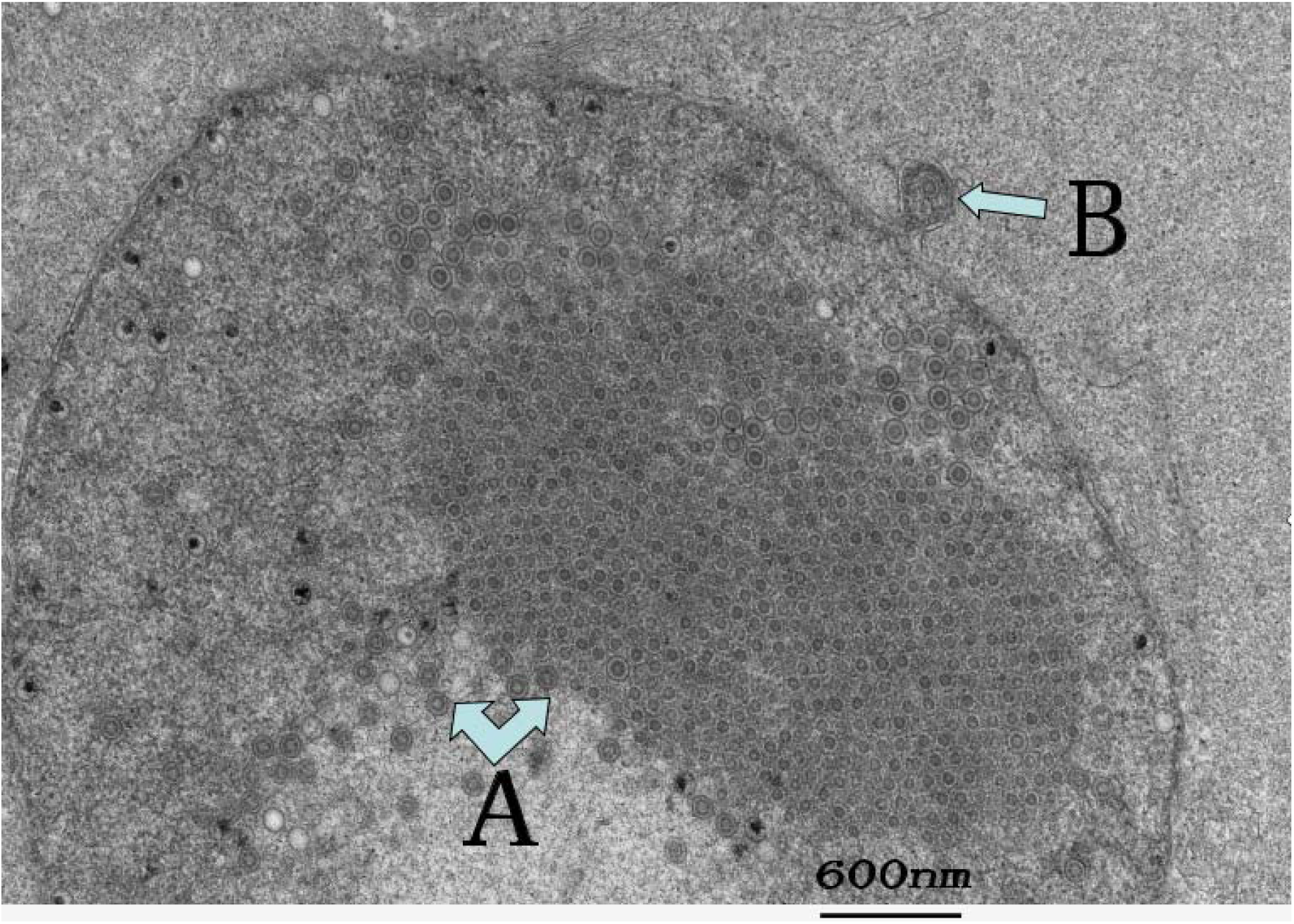
Electron micrographs of goldfish cells(GFB) infected with CyHV-2. A: large number of hexagonal nucleocapsid aggregates inside the nucleus;electron dense, double-layered ring shape and electron lucent nucleocapsid also observed in this aera; B: a virus inside the vesicles protruding outside the nuclear membrane;

**Figure 7.**
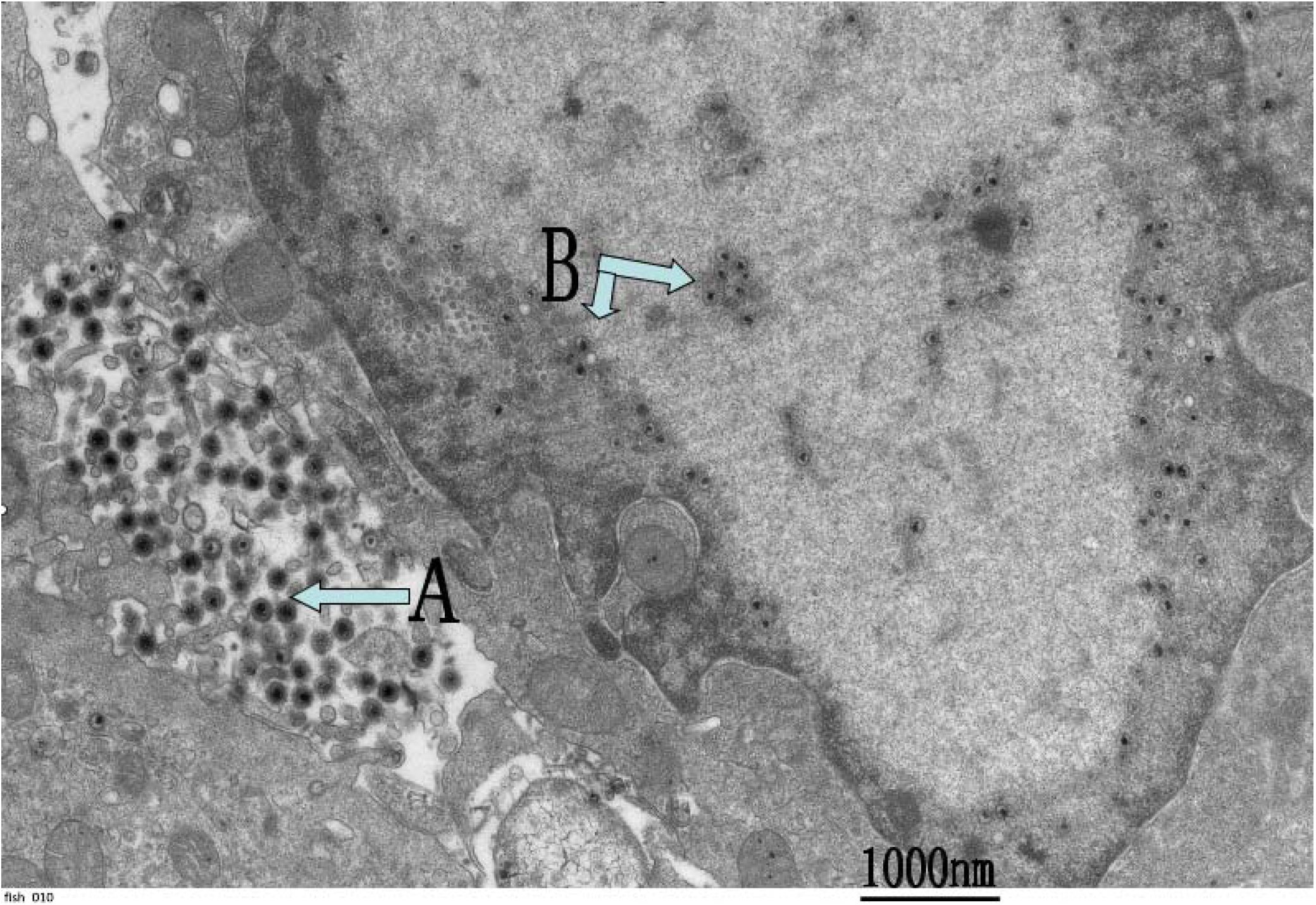
Electron micrographs of goldfish cells(GFB) infected with CyHV-2. A: A large number of electron dense spherical virus particles are present in the extracellular cytoplasm. Size about 200 nm. B: hexagonal nucleocapsid, about 110 nm in size, is visible in the nucleus.

Whole Genome analysis showed that 8 strains of CyHV-2 can be divided into 3 groups (Figure 8). Group 1 includes two ST-J1Strains (JQ815364.1 and NC019495.1) being 100% homology(identical) although they have different accession number. Group 2 includes strain SY. Group 3 includes the remain strains. YZ01 is closer to Cyprinid herpesvirus 2 strain CNDF-TB2015 and SY-C1 and belongs to same group. PhyML online execution showed that YZ01 was closer to YC-01 and belonged to a group of SY-C1, CNDF-TB2015, 1301, YC-01 and YZ01(data not shown), which was similar to those by MEGA X.

**Figure 8.**
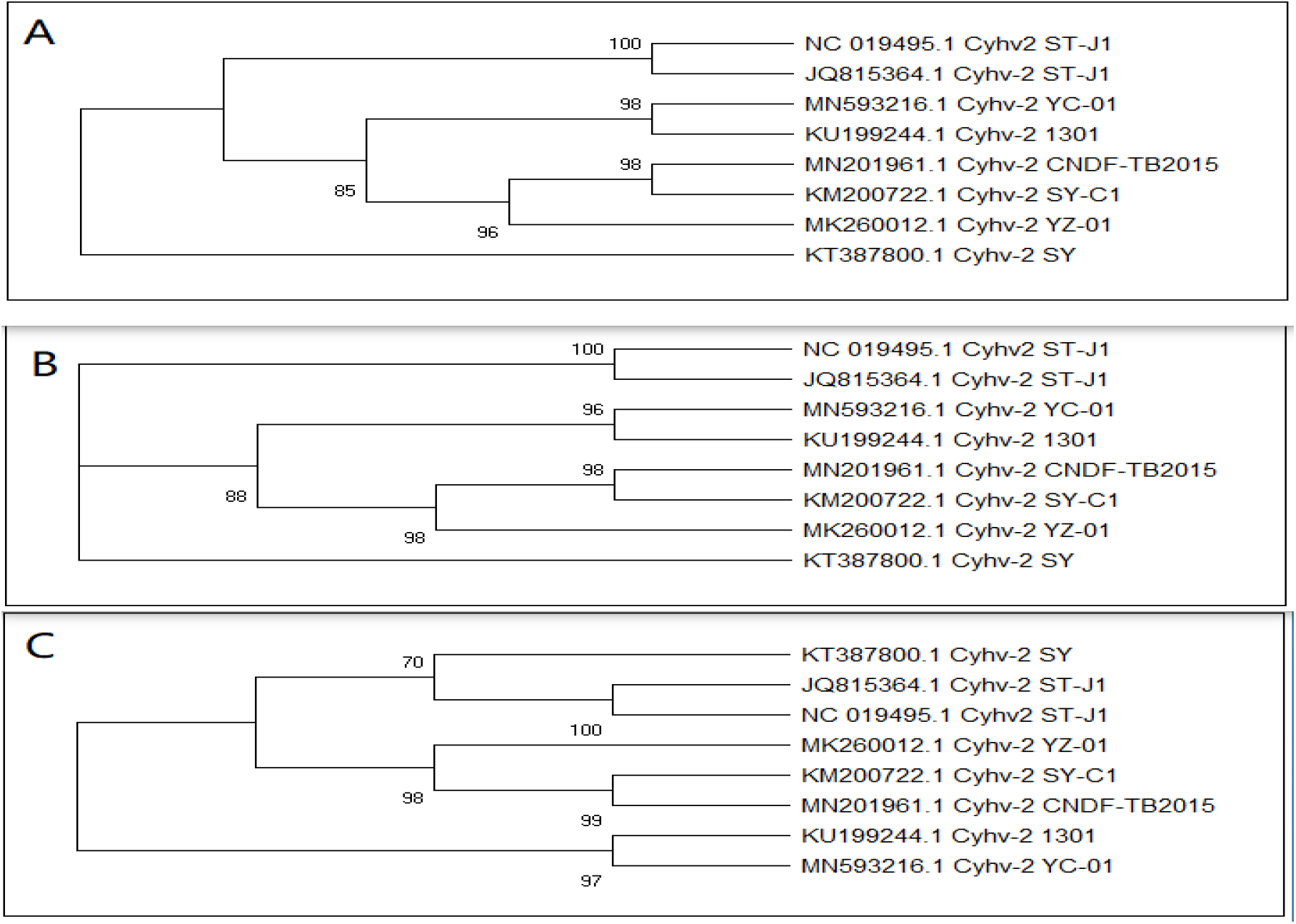
Evolutionary relationships of YZ01 and other 7 strains. The evolutionary history was inferred using: A-Minimum Evolution method(ME), B-the Neighbor-Joining method (NJ) and C-UPGMA method. The bootstrap consensus tree inferred from 1000 replicates is taken to represent the evolutionary history of the strains analyzed. The percentage of replicate trees in which the associated strains clustered together in the bootstrap test are shown next to the branches.

## Discussion

Research of fish herpesvirus suggested latent infection and/or carrier status persisted in clinically healthy fish and permissive temperature existed for the virus replication (Dishon et al., 2007; Gilad et al., 2004; Ito & Maeno, 2014; Nanjo et al., 2016; Uchii et al., 2013; Wei et al., 2019). In wild sero-positive carp replication-related genes of CyHV-3 were transcribed only under permissive water temperatures during the spring. While under non-permissive conditions possible latency-related genes were expressed and CyHV-3 did not multiply. The findings proposed that CyHV-3 established latent infection in persistent carriers and then reactivated periodically under increasing spring temperature (Uchii et al., 2013). In this challenge experiment, fish from a group with history of exposure to CyHV-2 began to die a week after the water temperature was raised from 3-6°C to about 25°C, and PCR results were positive for intestine and mixture samples of diseased or dead fish. 2-3 days after the PCR negative fish were transferred to the sick tank, the fish also started to die and were positive for PCR. This suggested that increased water temperature to permissive could reactivate virus replication in apparently healthy fish exposed to CyHV-2 earlier. It also indicated existence of horizontal transmission of the disease. Experimental infection of goldfish under different water temperature with CyHV-2 by intra-peritoneal injection and bath immersion showed 20-25°C was permissive temperature for HVHN(Ito & Maeno, 2014). Surviving fish in 15°C group and the fish co-habited with them died after temperature increased to 25°C. The above study also assumed that most fish infected with CyHV-2 at 13-15°C neither died nor acquired resistance to HVHN but able to infect naïve fish as carriers. CyHV-2 is often considered to be a latent virus taking account of the long incubation time of the disease and the fact that the disease is usually triggered by predisposing factors such as water temperature changes or stress(A. E. Goodwin et al., 2009). Other studies also found persistency status or latent infection of Cyprinid herpesvirus mainly CyHV-2 and CyHV-3 in fish or in cell lines (Imajoh et al., 2014; Reed et al., 2014; Wei et al., 2019). Viral load of CyHV-2 in an apparently healthy goldfish varied as temperature changed. A recent investigation of latency of CyHV-2 in survival fish after primary infection showed that CyHV-2 genomic DNA was detected in various tissues from fish of acute infection, but significantly reduced in fish recovered from the primary infection on 300 dpi(day post-infection). No active viral gene transcription was detected in recovered fish. After raising temperature, an increase of CyHV-2 DNA load and gene transcription were observed in tissues examined (Chai et al., 2020).

Development of cyto-pathological effects was very rapid and obvious after inoculation with infected CrCB or GFB cell suspension or tissue homogenate of dead fish. 4 days after inoculation of GFB cell line with brain, kidney and spleen (BKS) suspension from the dead goldfish of tank 4, cells in focal areas became brighter and round and intercellular space became clear and then focal lesions including cell damage forming network or sieve-like and cells aggregated into clusters(Figure 2).

Electron microscopy of goldfish cells (GFB) infected with tissue homogenate of dead fish with history of exposure to CyHV-2 in this study demonstrated cell lesion including degenerated or destroyed organelles, incomplete and invisible nucleus membranes. Large number and lattice-like arrangement of hexagonal nucleocapsid aggregate inside the nucleus was observed. Electron dense, double-layered ring shape and electron lucent nucleocapsids may be their precursor forms. CyHV-3 nucleocapsids were classified into three different types: capsids with an inner-loop (spherical) structure, electron-containing dense nucleocapsids, and empty capsids. These morphologies may represent three different developmental stages or phases of CyHV-3 capsid morphogenesis(Miwa et al., 2007). In this study, similar viral particles were observed by TEM in CyHV-2 infected GFB and CrCB cells. CyHV-2 and CyHV-3 have similar viral forms. However there were more viral particles observed in this experimental infected cells. Virus inside the vesicules protruding outside the nuclear membrane and vesicles containing virus in cytoplasm and a lot of electron dense spherical virus particles present outside of cells were also found. The size of nucleocapsids and virus particles, different stages and forms of virus in nucleus, cytoplasm and outside of cells of infected tissues were also reported in other studies((Thangaraj et al., 2020). These findings are in consistence with earlier researches on herpesvirus and support capsids replicate and assemble in the nucleus, mature capsids bud through the nucleus membrane and channel through endoplasmic reticulum and mature virions release outside of cells and infect adjacent cells. Aggregation of capsids and cell damage is in accordance with the rapid appearance of cell pathological effects observed in cell culture.

Whole genome analysis demonstrated that isolate YZ01 was closer to strain CNDF-TB2015 and SY-C1 and belonged to the same group (Figure 8). Cyprinid herpesvirus 2 strain CNDF-TB2015 was submitted on Jul 20, 2019 by Pan, X., Yuan,X., Lin, L. and Shen, J with tilte *Comparative genomics of two stains of Cyprinid herpesvirus 2 from China* (unpublished). This strain was isolated from spleen and kidney tissue of mature crucian carp *Carassius auratus* collected on Oct 8, 2015 in Dafeng, Yancheng, Jiangsu Province. A recent study showed that variant CyHV2-SY isolated from Sheyang, a county of Jiangsu Province shares 98.4% and 99.1% sequence identity with strains ST-J1 and SY-C1, respectively(Liu et al., 2018). That paper also proposed the significant genome diversity of CyHV2 strains isolated from different hosts was due to co-evolution with its own host and the host adaptation. Phylogenic tree analysis of the amino acid sequences of 12 core ORFs showed that strains SY, SY-C1 and CaHV from ACC were clustered together. Crucian carp *Carassius auratus* herpesvirus (CaHV) (Isolate 1301, accession number is KU199244)was isolated from crucian carp *Carassius auratus* in 2013 in China (NCBI). Phylogenic analysis of DNA polymerase genes of 17 herpesviruses and 12 concatenated core genes from known alloherpesviruses showed that CaHV was most closely related to SY-C1.But CaHV was proposed as a novel herpesvirus because of considerable genetic variations and lack of a direct repeat (Zeng et al., 2016). Dafeng(CNDF-TB2015) and Sheyang(SY) are in the same city Yancheng(YC) and a crucian carp aquaculture area and near where strain YZ01 was discovered. CyHV-2 strain SY (KT 387800.1) was formed a separate branch from other CyHV-2 strains by MEGA X although it was also isolated from spleen and kidney tissue from *Carassius auratus gibelio* suffering from hemopoietic necrosis in Jiangsu province. Further study is needed to investigate the genomic and molecular epidemiological characteristics of these CyHV-2 strains including CyHV-2 strain YZ01.

## Conclusion

CyHV-2 strain YZ01 was isolated from apparently healthy goldfish after rising water temperature. Electron microscopic examination of the YZ01 infected GFB cell line and whole genome comparison published in GeneBank were performed to further confirm that the isolate had the characteristics and epidemiological profile of Cyprinid herpesvirus 2 of Cyprinidae.

## Competing interests

The authors declare that they have no competing interests.

## Acknowledgments

This work was financially supported by the science and technology project of Nanjing Customs of P.R. China (No.2019KJ16) and Jiangsu Entry-Exit Inspection and Quarantine Bureau (No. 2017KJ01).

## Notes

### Competing Interest Statement

The authors have declared no competing interest.

